# Mitochondrial function regulates cell growth kinetics to actively maintain mitochondrial homeostasis

**DOI:** 10.1101/2025.03.31.646474

**Authors:** Leeba Ann Chacko, Hidenori Nakaoka, Richard Morris, Wallace Marshall, Vaishnavi Ananthanarayanan

## Abstract

Mitochondria are not produced *de novo* in newly divided daughter cells, but are inherited from the mother cell during mitosis. While mitochondrial homeostasis is crucial for living cells, the feedback responses that maintain mitochondrial volume across generations of dividing cells remain elusive. Here, using a microfluidic yeast ‘mother machine’, we tracked several generations of fission yeast cells and observed that cell size and mitochondrial volume grew exponentially during the cell cycle. We discovered that while mitochondrial homeostasis relied on the ‘sizer’ mechanism of cell size maintenance, mitochondrial function was a critical determinant of the timing of cell division: cells born with lower than average amounts of mitochondria grew slower and thus added more mitochondria before they divided. Thus, mitochondrial addition during the cell cycle was tailored to the volume of mitochondria at birth, such that all cells ultimately contained the same mitochondrial volume at cell division. Quantitative modelling and experiments with mitochondrial DNA-deficient *rho0* cells additionally revealed that mitochondrial function was essential for driving the exponential growth of cells. Taken together, we demonstrate a central role for mitochondrial activity in dictating cellular growth rates and ensuring mitochondrial volume homeostasis.

## Introduction

As a cell grows, the size or volume of organelles such as the nucleus, lysosomes and mitochondria typically scales with cell size (1–4). This scaling relationship is attributed to the increasing functional demands on the organelle as the cell grows. However, the mechanisms that underpin organelle scaling and homeostasis are poorly understood. For example, while the importance of mitochondrial activity for cellular fitness is widely accepted (5–8), it remains unclear how mitochondrial volume is maintained across successive generations.

Mitochondria typically form tubular networks that undergo continuous cycles of fusion and fission, enabling the redistribution of mitochondrial DNA (mtDNA) and the dilution or removal of damaged components (9–11). Since mitochondria are composed of proteins that are encoded by both nuclear and mtDNA, mitochondria cannot be synthesised *de novo* in newly-divided daughter cells, but are partitioned into daughter cells from the mother. In fission yeast, we recently described an independent segregation mechanism for mitochondrial partitioning (12). Mitochondria in mother cells undergo enhanced fission upon loss of microtubule association during closed mitosis in fission yeast, which increases mitochondrial copy numbers and thus reduces partitioning errors in this system (12). Regardless, due to the nature of the process, mitochondrial partitioning exhibited binomial errors (13), resulting in slight differences in mitochondrial volumes at birth even in these symmetrically dividing cells.

Cell size homeostasis in fission yeast is achieved through the ‘sizer’ mechanism, wherein cells sense their own size and a critical size threshold is reached before cells undergo division (14–16). Using mutants with varying widths compared to WT cells, fission yeast has been found to undergo division upon reaching a defined surface area, in a process dependent on the protein kinase Cdr2 (16–18). Other systems, including bacteria and mammalian cells behave as ‘adders’, adding a constant mass during each cycle (19, 20).

Recently, we discovered that fission yeast cells with mutations or deletions of proteins responsible for microtubule and mitochondrial dynamics exhibited a significant degree of asymmetric cell division (21). Asymmetrically-dividing cells still followed independent segregation, with the smaller daughter cell receiving less mitochondria from the mother than the larger daughter cell. Occasionally, we observed mutant cells that partitioned their cytoplasm symmetrically but their mitochondria asymmetrically; in these cases, the daughter cell with less mitochondria grew slower than the other daughter. Given this key role of mitochondrial function in dictating cell growth and fitness, we asked how mitochondrial homeostasis is maintained across several generations of dividing cells.

Here, we use extended live cell imaging in a yeast mother machine (YMM) (22, 23), automated analysis, and quantitative modeling to demonstrate the critical role of mitochondrial function in determining not only feedback-driven mitochondrial homeostasis, but also the timing of cell division, which ultimately actively controls cell size homeostasis in fission yeast.

## Results

### Fission yeast cells achieve cell size homeostasis through the sizer mechanism

To visualise growth and division in several generations of fission yeast cells, we employed a YMM (Fig. S1A). We confirmed that cells were healthy in the YMM, since the cell dimensions over the course of their cell cycle (Fig. S1B, Supplementary Video S1) were consistent with published literature (24–28). Similarly, the cell lengths at birth (8.1 ± 0.8 *µ*m; mean ± standard deviation (SD), n=2159 cells, see Supplementary Table S1) and at division (15.8 ± 1.4 *µ*m; mean ± SD, n=2159 cells) were comparable to the known estimates in fission yeast cells (24–27), and so was the mean duration of a cell cycle at 1.8 ± 0.3 h (mean ± SD, n=2159 cells, Fig. S1C-D).

We then probed the size homeostasis mechanism in wildtype (WT) cells growing in the YMM and reconfirmed several previous findings suggesting that fission yeast cells show sizer-like behaviour (14, 16, 24, 29), with cells that were smaller than the average length of ~8.1 *µ*m at birth adding more length before division and vice versa (Fig. S2A-C), ultimately resulting in cells initiating division at a characteristic size of 15.8 ± 1.4 *µ*m (mean ± SD, n=2159, Fig. S2D). Another consequence of the sizer-like behaviour of fission yeast was that cells that were born smaller than average exhibited longer cell cycle durations (Fig. S2E), as reported previously (30, 31). We additionally confirmed that cellular volume, surface area and length reported similarly on the cell size for the duration of the cell cycle (Fig. S3A).

We previously showed that fission yeast cells with mutations or deletions of mitochondrial morphology-related proteins exhibit a higher degree of asymmetric cell division compared to WT cells (21). Cells devoid of microtubule-associated proteins are also known to exhibit asymmetric cell division (12, 32). We asked if these mutants that exhibit a high degree of asymmetric cell division also showed sizer-like behaviour for cell size homeostasis. We observed that similar to WT cells, these mutant cells tailored their cell size at division to their size at birth - cells that were longer than average at birth divided faster in the following cycle to compensate for increased size at birth (Fig. 1A-C, Supplementary Video S2). These asymmetrically-dividing mutants also showed a characteristic division length, confirming their sizer-like behaviour for size homeostasis (Fig. 1D, see Supplementary Table S1).

**Figure 1.**
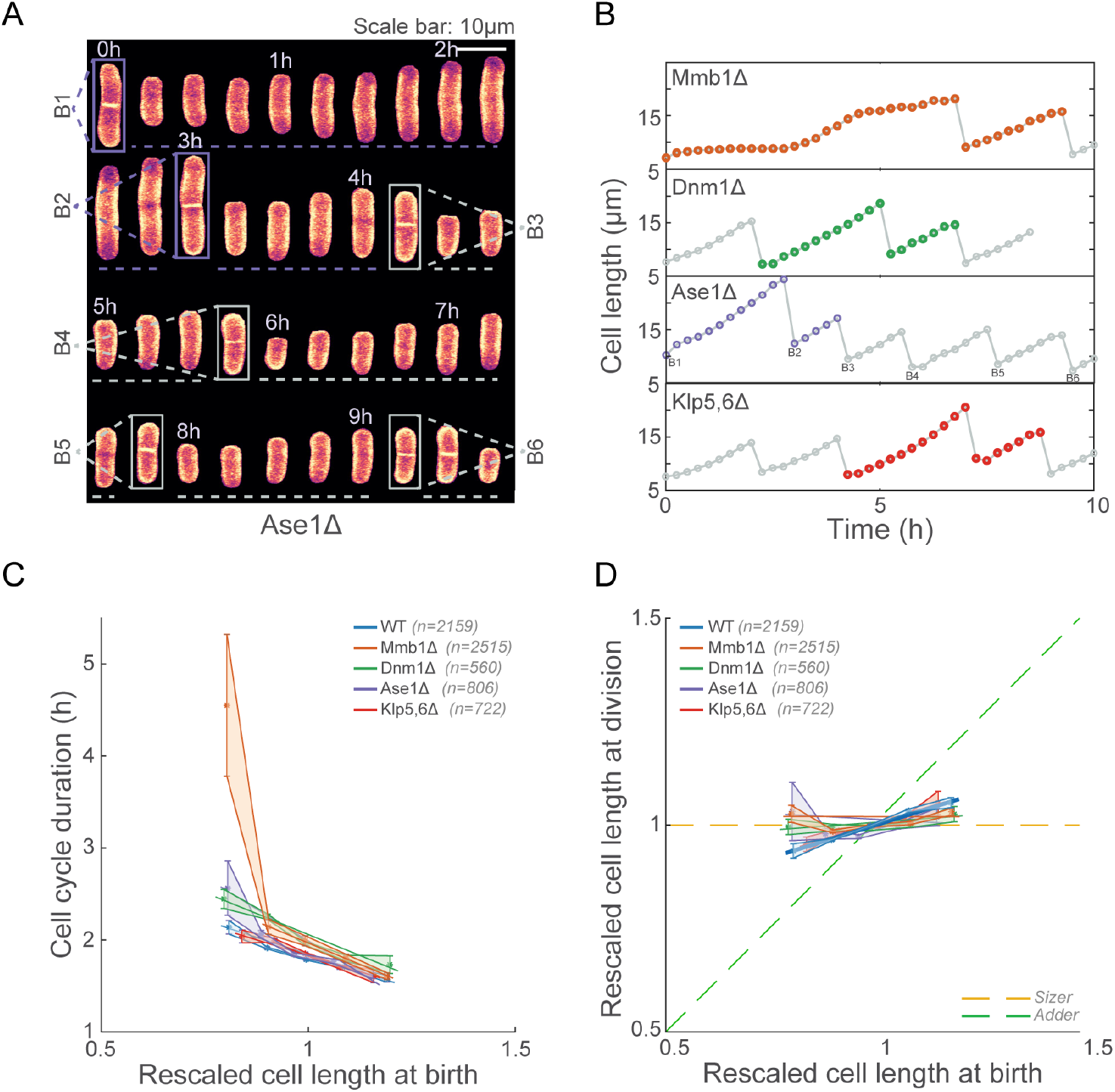
Mutants with a high degree of asymmetric division also exhibit “sizer-like” behaviour. **(A)** Representative montage of an Ase1Δ cell (expressing a fluorescent membrane marker) undergoing growth and division in a single channel of the yeast mother machine (YMM). **(B)** Plots of cell length over time for all the mutants indicated. Coloured circles represent a long cell cycle followed by a short cycle. ‘B1’-’B6’ in the Ase1Δ plot represent 6 individual cell birth events as indicated in **A. (C)** Cell cycle duration is inversely proportional to the cell size at birth, as shown by the cell cycle duration plotted against the cell length at birth in wild-type (WT) and mutant cells. Note that the Mmb1Δ fit was computed using only the last four bins, as the first bin contained several outliers. **(D)** Cell length at division plotted against cell length at birth in wild-type (WT) and mutant cells demonstrates that the sizer behaviour is active in these cells. The yellow dashed line represents the expected data for a perfect sizer and the green dashed line that for an adder. The plots in **C** and **D** show mean ± standard error of the mean (SEM) (solid circles and the shaded region respectively) and their fits (solid lines). The data in **C** and **D** were rescaled as detailed in the Methods section.The slopes and *R*^2^ values of the fits in **C** and **D** are in Supplementary Table S2. Strains VA130, VA131, VA136, VA135, and VA144 were used for all figures in this panel (see Supplementary Table S3).

### Mitochondrial volume is proportional to cell size

We then probed mitochondrial volume in WT and mutant cells for the duration of their cell cycle (Supplementary Video S3). We previously demonstrated that mitochondrial partitioning at division proceeds relative to daughter cell sizes (21). Here, we tracked the mitochondrial volume alongside the cell sizes (Fig. S3A) and observed that mitochondrial volume at birth scaled with the cell size at birth, such that a smaller cell than average contained less mitochondria at birth than a larger cell (Fig. 2A-B). Mutant cells with higher degrees of asymmetric cell division also exhibited similar behaviour (Fig. S3B), with the mitochondrial volume inherited during division showing a linear relationship to cell size at birth (Fig. 2C).

**Figure 2.**
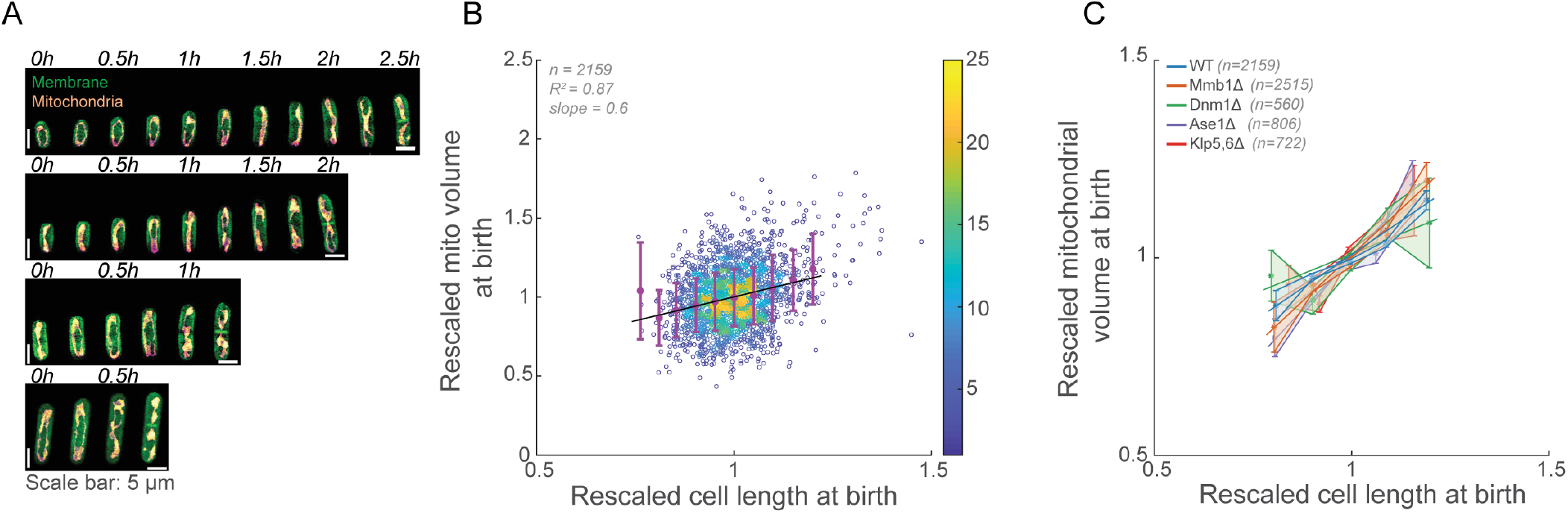
Mitochondrial volume at birth scales with cell length at birth. **(A)** Example montages of cell cycles of different sized WT cells expressing fluorescent membrane and mitochondria markers, **(B)** Mitochondrial volume at birth plotted against cell length at birth in WT cells reveals a linear scaling between the two parameters. Error bars represent SD and the black solid line represents a linear fit to the data; the heat map represents the density of the data. **(C)** Mitochondrial volume at birth plotted against cell length at birth in mutant cells also demonstrates a similar linear relationship between cell size and mitochondrial volume at birth. The plots show mean *±*^2^ SEM (solid circles and the shaded region respectively) and their fits (solid lines). The data in **B** and **C** were rescaled as detailed in the Methods section. The slopes and *R* **C** are in Supplementary Table S2. Strains VA130, VA131, VA136, VA135, and VA144 were used for this panel (see Supplementary Table S3).

### Mitochondrial volume at division exhibits compensatory signatures

Next, we asked how mitochondrial homeostasis was maintained across several generations. A previous study in fission yeast suggested that a constant volume of mitochondria is added during each cycle irrespective of the mitochondrial volume at birth (33). Alongside our data suggesting that mitochondrial volume at birth is dependent on cell size at birth (Fig. 2), this would imply that even a cell that received less volume at birth (due to its smaller size) would add the same volume of mitochondria as a large cell. We measured the mitochondrial volume added during a cycle (by comparing the mitochondrial volume at division and at birth), and plotted this against the cell length at birth (Fig. 3A). We observed that contrary to the proposed mechanism of constant volume addition, cells that were smaller than average at birth added more mitochondrial volume during their cell cycle. This resulted in cells, small and large, achieving similar mitochondrial volumes at division (Fig. 3B), suggesting that rather than adding constant volume, cells exhibited compensation of mitochondrial volumes by adding more mitochondrial volume when born with less, and vice versa. So too, the volume added per unit length before cell division was similar for cells with a wide range of starting mitochondrial concentrations (mitochondrial volume per unit length at birth) (Fig. 3C), i.e., cells that were small at birth added more length (Fig. S2C) *and* mitochondrial volume such that the ratio between the added mitochondrial volume and added cell length remained a constant. We previously showed that cells that were smaller than average at birth had longer cell cycle durations (Fig. S2E). Accordingly, cells that had longer cell cycle durations also added more mitochondria than those that had shorter cell cycle durations (Fig. S4A), which fits with the compensatory homeostatic responses we observed.

**Figure 3.**
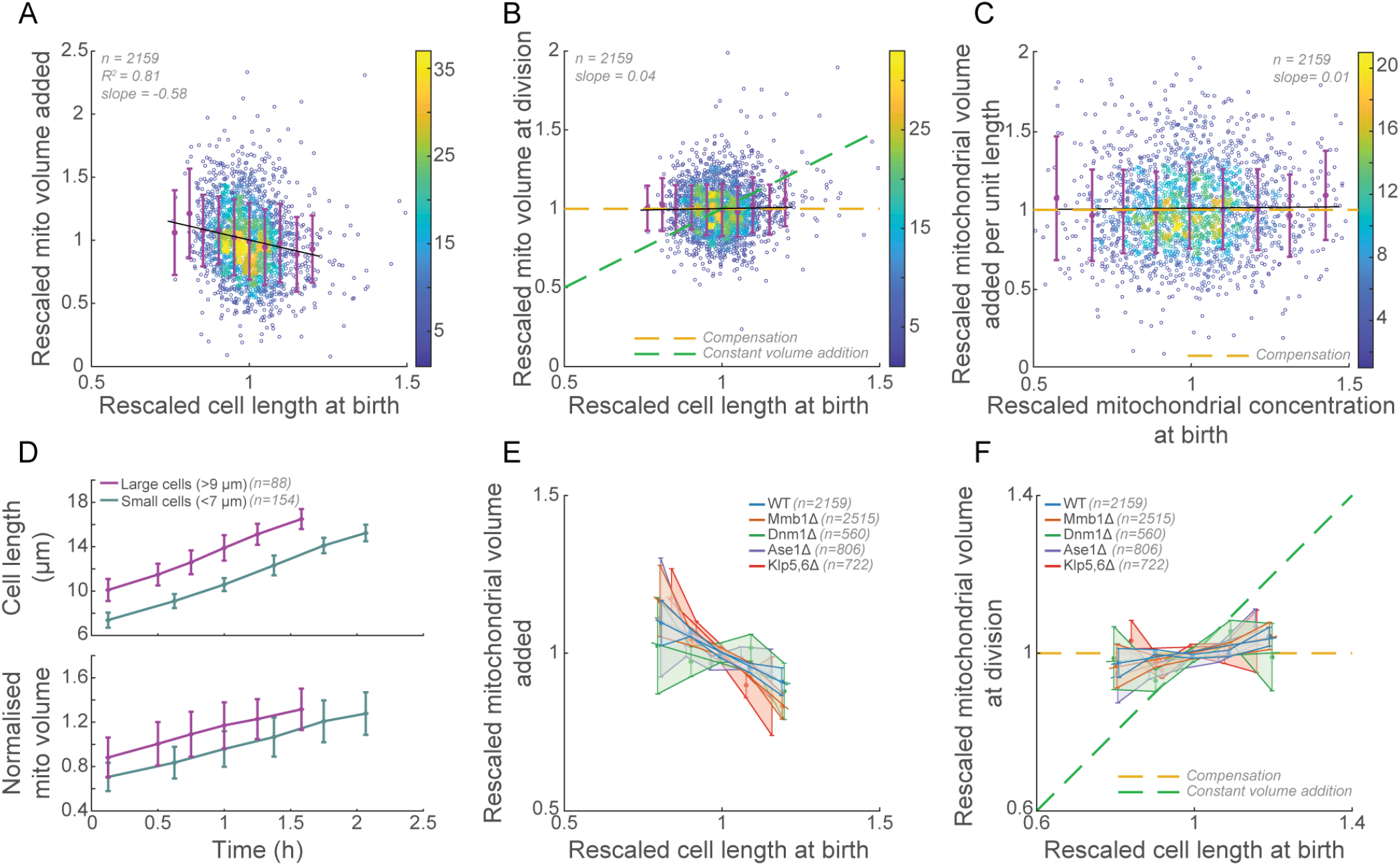
WT and mutant cells exhibit ‘compensation’ of mitochondrial volume to achieve homeostasis. **(A)** Mitochondrial volume added in a cycle plotted against the cell length at birth in WT cells reveals an inverse relationship. **(B)** Mitochondrial volume at division plotted against the cell length at birth in WT cells demonstrates signatures of compensation, wherein all cells attain a typical mitochondrial volume at division. **(C)** Mitochondrial volume added per unit length added during the cell cycle plotted against the mitochondrial concentration at birth in WT cells reveals a compensatory mechanism, with a constant volume of mitochondria added per unit length of cell. In **A-C**, error bars represent SD. Black solid lines represent linear fits to the data; the heat map represents the density of the data. **(D)** Cell length (top) and the normalised mitochondrial volume (bottom) plotted over time in large (>9 *µ*m, magenta) and small (<7 *µ*m, teal) WT cells reveals differences in the duration of the cell cycle between the two populations. Error bars represent SD. **(E)** Mitochondrial volumes added per cell cycle plotted against cell lengths at birth in mutant cells also shows an inverse relationship. **(F)** Mitochondrial volumes at division plotted against cell lengths at birth in mutant cells also demonstrates mitochondrial compensation. In **B, C** and **F**, the yellow dashed line represents the expected data for a compensation mechanism and the green dashed that for constant volume addition. The plots in E and F show mean^2^*±* SEM (solid circles and the shaded region respectively) and their fits (solid lines). The data in **A**-**F** were rescaled as detailed in the Methods section. The slopes and *R* values of the fits in **E** and **F** are in Supplementary Table S2. Strain VA130 was used for panels **A-D** and strains VA130, VA131, VA136, VA135, and VA144 were used in panels **E** and **F** (see Supplementary Table S3).

Further, we split the WT cells into two populations - large and small cells, with the former having cell lengths >9 *µ*m and the latter with cell lengths <7 *µ*m at birth, i.e., 1 SD away from the mean length at birth. We tracked their cell lengths and mitochondrial volumes over the course of the cell cycle and observed that while the small cells started with lower cell lengths and mitochondrial volumes, they attained similar lengths and mitochondrial volumes as the large cells at the end of their cycle by prolonging their cell cycle (Fig. 3D). Thus, fission yeast cells rely on the cell sizer mechanism to maintain mitochondrial homeostasis: larger *(smaller)* cells typically contain more *(less)* mitochondria at birth, but need to add less *(more)* cell size (length/area) before division, thus reducing *(increasing)* the amount of mitochondria added during the cell cycle. We observed similar behaviour in mutant cells that have a higher propensity to divide asymmetrically, which showed an inverse relationship between cell length at birth and mitochondrial volume added during the cell cycle (Fig. 3E), resulting in a characteristic mitochondrial volume at division across the population (Fig. 3F). Thus, both WT and mutant cells corrected for deficiencies or excess mitochondrial volume in one generation, as indicated by the similar distributions of mitochondrial volumes at birth for these cells (Fig. S4B-F).

### Mitochondrial activity drives the exponential growth of cells and determines the timing of cell division

Thus far, we determined that mitochondrial homeostasis relies on the cell sizer mechanism. However, given the dependence of starting mitochondrial volume on cell size at birth, it remained unclear what the precise role of the sizer was in determining mitochondrial homeostasis. To delineate the specific role of mitochondrial activity, we chose to analyse daughter cell pairs in the WT population that divided symmetrically, i.e., had similar cytoplasmic volumes, but exhibited asymmetric mitochondrial partitioning. Such daughter cell pairs arose due to the nature of mitochondrial partitioning in fission yeast that follows an independent segregation mechanism, with binomial errors (12). Analysis of the cell lengths over time (Fig. 4A) of these daughter cell pairs revealed that the cell with more mitochondria at birth grew significantly faster than the other daughter cell (Fig. 4B), thus reaching the characteristic cell length for the sizer mechanism to operate sooner than the other daughter cell (Fig. 4C). We confirmed that the daughters with more and less mitochondria still exhibited their sizer-like behaviour by comparing the cell lengths at division, and observed that both populations did indeed achieve similar lengths at division (Fig. 4D top). So too, cells born with different starting mitochondrial volumes showed a similar mitochondrial volume at division (Fig. 4D bottom), indicating the presence of compensatory mechanisms. Together, these data suggest that mitochondrial volume at birth is a key determinant of the timing of cell division in fission yeast.

**Figure 4.**
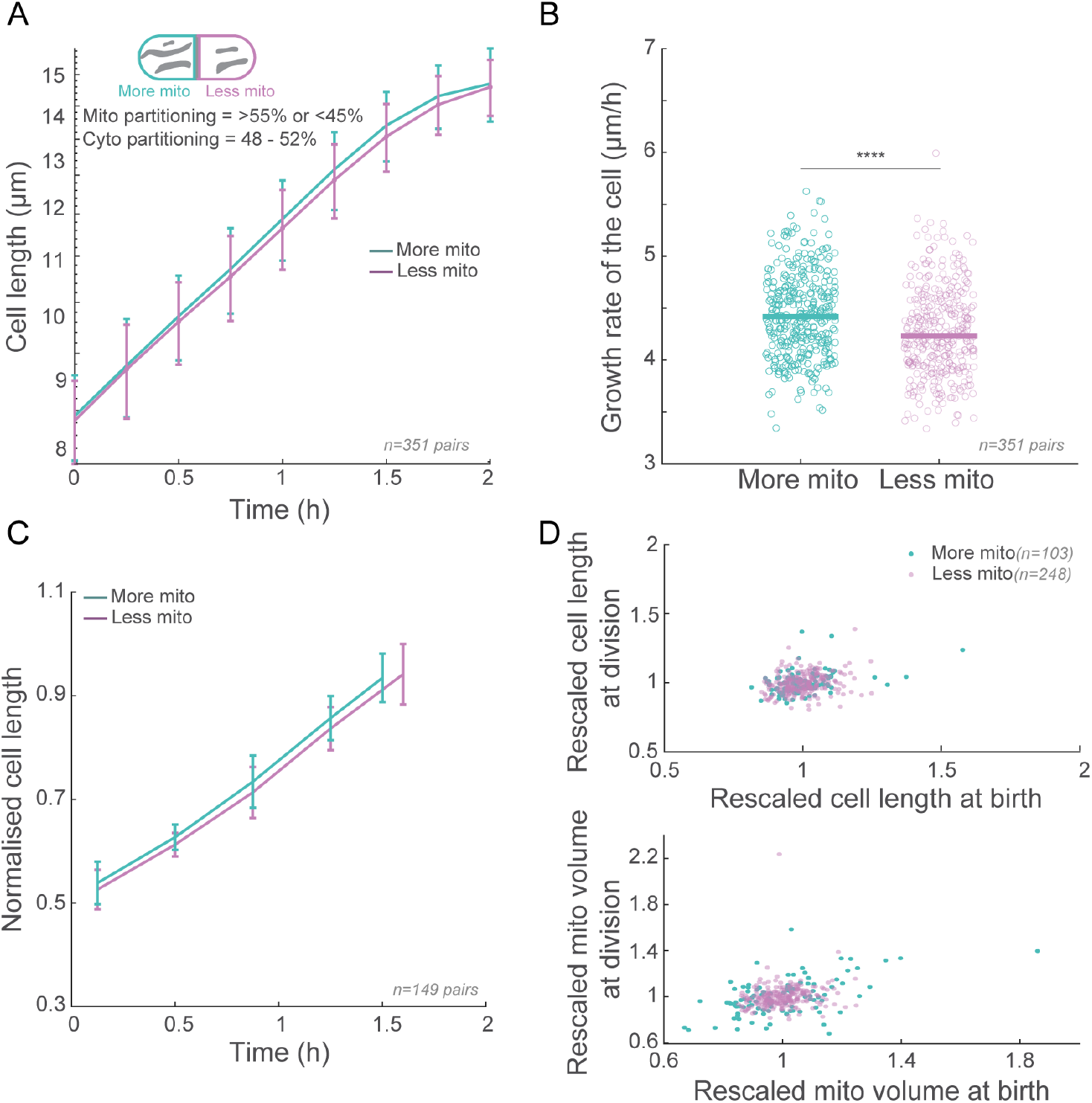
Mitochondrial activity determines the timing of cell division. **(A)** Cell length plotted over time for pairs of daughter cells with similar cell sizes, but unequal mitochondrial volumes (‘More mito’ and ‘Less mito’, with >55% of the mother cell mitochondrial volume partitioned in the former, and <45% partitioned into the latter, but with both daughters having 48-52% of the mother cell length at division). The solid line represents the mean and the error bars represent SD. **(B)** Scatter plot of the growth rate of the cells analysed in **A**. The horizontal lines represent the mean. The asterisks represent significance (**** = *p <* 10^*−*4^); unpaired T-test for parametric data. **(C)** Normalised cell length plotted over time for those pairs of cells whose cycles were tracked fully indicating a longer cell cycle duration for those cells that had less mitochondrial volume at birth. The solid line represents the mean and the error bars represent the SD. **(D)** Top - scatter plot of cell length at division versus cell length at birth for the data in **C** showing that cells still behave as sizers regardless of starting mitochondrial volume. Bottom - scatter plot of mitochondrial volume at division versus that at birth for the data in **C** demonstrating mitochondrial compensation in cells born with more and less mitochondria than average.The magenta circles represent cells in the daughter pair with less mitochondria (<45% mitochondrial volume partitioned from the mother) and the teal circles represent cells with more mitochondria (>55% mitochondrial volume partitioned from the mother). Strain VA130 was used for all panels in this figure (see Supplementary Table S3).

To further explore the role of mitochondria in determining the growth kinetics and hence the timing of cell division, we tracked the change in cell size and mitochondrial volume over time in WT cells (Fig. 5A). Measurement of the instantaneous growth rates of cells and the size-normalised instantaneous growth rates over time reconfirmed earlier findings that cells grow exponentially until mitosis in fission yeast (Fig. 5B, S5A). Thereafter, the cellular growth rate slowed down considerably until cells eventually divided. However, unlike cellular growth, mitochondrial addition rate continued to increase, implying an exponential increase in mitochondrial volume throughout the cell cycle (Fig. 5B, S5B). Given our finding that in a daughter cell pair, the daughter that received more mitochondria from the mother grew faster (Fig. 4B), we probed if the exponential growth of cells was driven by the exponential addition of mitochondria and the consequent exponential increase in mitochondrial activity. We modeled mitochondrial volume to increase exponentially over time. Specifically, we describe mitochondrial activity *M* as:

**Figure 5.**
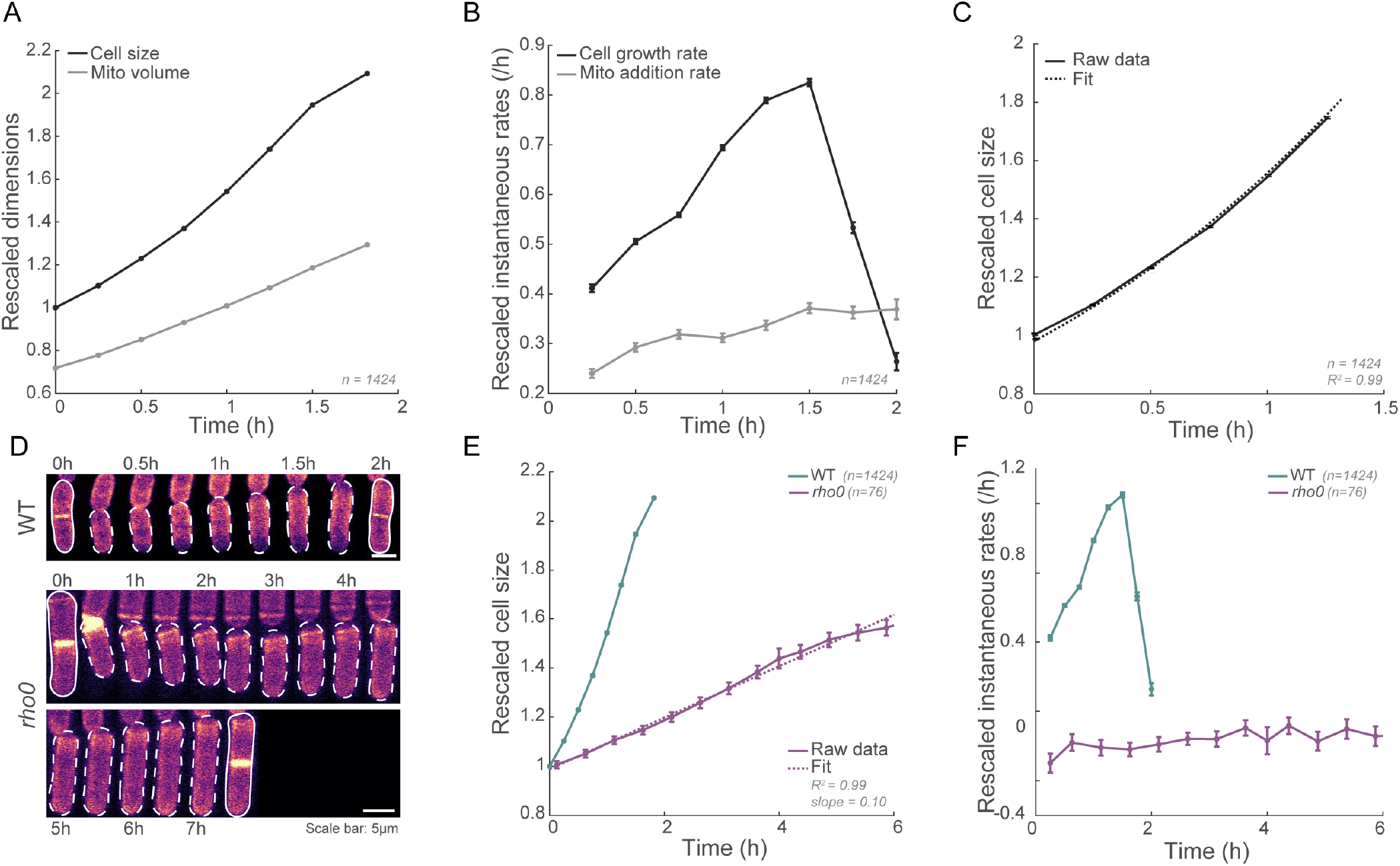
Loss of mitochondrial activity leads to linear cell growth. **(A)** Plots of cell size (area, black) and mitochondrial volume (grey) over time for those cells whose durations are within one SD of the mean WT cell cycle duration. Error bars represent SEM. **(B)** Plots of instantaneous growth rates of the cell (black) and mitochondrial volume (grey) respectively for the cells analysed in **A**. Error bars represent SEM. **(C)** Plot of cell area *A* over cell cycle duration *t* (solid black line, excluding the M-phase). The dashed black line represents a fit to the equation *A* = 2.57 *e*^0.32*t*^ *−* 0.39*t* + *E*. Error bars represent standard error of the mean. **(D)** Representative montages of a WT cell (top, expressing a fluorescent membrane marker), and *rho0* cell (bottom, stained with calcofluor white) undergoing one cycle of growth and division. **(E)** Plots of cell size over time for WT and *rho0* cells (teal and magenta respectively) whose durations are within one SD of the mean WT and *rho0* cell cycle durations respectively. The dashed magenta line represents a fit to a polynomial with degree 1. Error bars represent SEM. **(F)** Plots of instantaneous growth rates for WT and *rho0* cells (teal and magenta respectively) for the cells analysed in **D**. Error bars represent standard error of the mean. The data in **A**-**C** and **E**,**F** were rescaled as detailed in the Methods section. Strain VA130 was used for panels **A-D** and strains VA130 and PHP14 were used for panels **D-F** (see Supplementary Table S3).

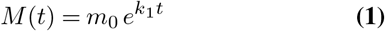

*M* is equivalent to mitochondrial volume in WT cells, *m*_0_ is mitochondrial volume at the start of the cell cycle, *t* is time and *k*_1_ is the mitochondrial volume addition rate.

The resulting fit to the raw data was highly robust, with an *R*^2^ value of 0.99 (Fig. S5C), confirming that an exponential model accurately captures the observed mitochondrial growth dynamics. From this fit, we extracted the values of *m*0 and *k*_1_.

Next, we modeled the growth rate of the cell 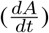 as dependent on the activity of the mitochondria:

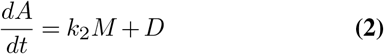

where *k*_2_ is a rate constant that links mitochondrial activity to cell growth and *D*, the mitochondrial activity-independent growth rate.

We plotted the cell growth rate 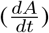 against mitochondrial volume (activity) (*M*) during the cell cycle, excluding the M-phase where cell growth stalled. We then performed a linear regression on the data to determine the value of *k*_2_. The linear fit achieved an *R*^2^ value of 0.97 ((Fig. S5D).

Finally, substituting (1) and the fitted values of *c* = 0.72, *k*_1_ = 0.32, *k*_2_ = 1.14 and *D* = −0.39 into (2):

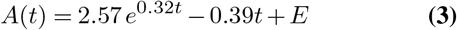

where *E* is the integration constant. Note that D does not accurately capture mitochondrial activity-independent growth rates in WT cells, since we do not have data points around M=0, i.e., cells whose starting mitochondrial volume/activity is close to zero. The value of D here is based on the fit to the available dataset, and this parameter is better estimated using a strain that lacks mitochondrial ATP production, as discussed below.

Fitting (3) to the plot of cell size *A* over time *t* (Fig. 5C) resulted in an *R*^2^ value of 0.995, indicating an excellent correlation between the model and the experimental data. This confirms that our model is sufficient to describe the dependence of the exponential growth of the cell on mitochondrial activity. This model also predicts that in the absence of mitochondrial volume or activity i.e., M=0, rather than exhibit exponential cell growth, cells would switch to linear growth, with a constant growth rate across the cell cycle. In other words:

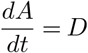

and

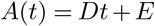

To test this prediction, we employed fission yeast cells lacking mtDNA (*rho0*), which do not generate mitochondrial activity-derived ATP, and hence are known to grow much slower than WT cells (34, 35). We compared the cell lengths over time and growth rates of WT and *rho0* cells (Fig. 5D) and measured a growth rate of only 1.1 ± 0.4 *µ*m/h (mean± SD, n=128 cells) for the *rho0* cells as opposed to 3.8 ± 0.4 *µ*m/h (mean ± SD, n=2159) in WT cells. Notably, while cell size increased exponentially until mitosis in WT cells, the growth rate was constant throughout the cell cycle in *rho0* cells as predicted by the quantitative model (Fig. 5E, F). So too, we observed a small proportion of one of our mutants (lacking the microtubule-mitochondria linker protein, Mmb1) partitioned mitochondria between daugther cell pairs substantially asymmetrically while still dividing symmetrically (also characterised in our previous work (21)). We observed the growth rate of the daughter cell that received very little mitochondria remained nearly constant throughout the cell cycle, while that of the other daughter cell increased during the first half (Fig. S5E, F) similar to WT cells, reiterating the critical role of mitochondrial volume/activity in determining cellular growth kinetics.

## Discussion

In this work, we used a YMM to observe several generations of fission yeast cells undergoing growth and division (Fig. S1). We first confirmed that both WT cells and mutant cells with a propensity for asymmetric cell division displayed sizer-like behaviour, undergoing division at a characteristic size (Figs. S2, 1). While the sizer property of WT cells has been well-established (16–18, 24, 36), that mutant cells demonstrate similar behaviour is interesting, and points to a decoupling of the machinery involved in establishing the cell division plane and that responsible for enforcing the sizer. We additionally observed that the cell cycle duration is inversely proportional to cell size at birth (Fig. 1C), as described previously (24).

We quantified mitochondrial volume of cells from birth to division and discovered that while mitochondrial volume partitioning scaled with the birth size of the cell (cells that are small at birth typically have less mitochondria than large cells, Fig. 2B, C), all cells attained similar mitochondrial volumes at division (Fig. 3A, B). This finding reveals a novel compensatory mechanism that corrects for initial mitochondrial “deficits” or “excesses”. In an earlier study in *S. pombe* (33), the authors concluded that mitochondrial volume homeostasis behaved like an adder wherein a constant volume of mitochondria was added per cycle regardless of the starting mitochondrial volume. However, a deeper look at their data suggests that the authors may have in fact measured mitochondrial compensation similar to our data, since mitochondria volume (intensity) added per unit length of growth during the cell cycle remained constant in their study (see Fig. 4C, in ref. (33), equivalent to 3C in this manuscript). This would imply that a cell that added more length during the cycle (i.e., one that was short at birth) would also add more mitochondrial volume, which is in agreement with our data (Fig. 3A, C). So too, if cells only added a constant volume of mitochondria per cycle, any deviation from the average mitochondrial volume at birth should take a few generations to converge back to the average volumes, as has been described for bacterial size regulation (19). In contrast, we found that deviation from mean mitochondrial volumes at birth were rectified within a single generation (Fig. 3B, S4B-F), which provides additional evidence of a compensatory mitochondrial homeostasis mechanism in these cells. Live-cell observations in *S. cerevisiae* have also revealed signatures of compensatory homeostasis, wherein a mother cell entering G1 with an abnormal mitochondrial volume relative to the total cell volume compensates by altering mitochondrial biogenesis to restore the correct ratio in the bud (3).

Given the tight coupling in cells between cell size and mitochondrial volume, we sought to disentangle their specific roles in the process of mitochondrial homeostasis. We previously described mitochondrial volume as a critical determinant of cell growth rates (21), but did not follow the fates of cells with altered mitochondrial volumes at birth. Here, we discovered that in daughter cell pairs that partitioned their cytoplasm symmetrically but exhibited asymmetry in mitochondrial partitioning, the daughter cell with reduced mitochondrial volume showed slower growth rates (Fig. 4). Other studies in mammalian cells report on similar phenomena, with daughter cells inheriting more mitochondria following cell division progressing faster through the cell cycle (37). Here, we found that while cells with less (*more*) mitochondria grew slower (*faster*), all cells exhibited sizer-like behaviour, dividing at similar sizes (Fig. 4C, D). Thus, mitochondrial compensatory mechanisms determine the time at which the sizer is enforced, but the molecules and processes controlling the sizer likely operate independently of the mitochondrial volume at birth.

*S. pombe* growth rates have been described as linear, bilinear, exponential, and non-exponential (15, 25, 28, 38–41). With the high-throughput dataset we acquired, we found that the growth of cells fit best to exponential growth kinetics until cells entered mitosis (Fig. 5B, S5A). We observed that mitochondrial volume also increased exponentially throughout the cell cycle (Fig. S5B). Thus, mitochondrial volume feeds back to the biogenesis machinery with genes encoded by both nuclear and mtDNA to enable exponential growth. A recent study in *S. pombe* compared the proteomes of large (*cdc25-22*) and small (*wee1-50*) mutants and found that mitochondrial proteins “super-scaled” with cell size, suggesting that bigger cells allocate disproportionately more resources to mitochondria (42). This observation contrasts with the idea that mitochondrial volume addition is a fixed or constant increment (33). Instead, the data indicate that as cells become larger, they likely ramp up mitochondrial biogenesis, potentially to meet the heightened metabolic and energy demands of a larger cytoplasm (42).

We found a key difference between cellular growth rate and mitochondrial addition rate in cells: while cellular growth rates declined and eventually almost completely stalled during the later stages of a cell cycle, mitochondrial volume continued to add at an increasing rate for the entire cycle (Fig. 5B). Such increases in protein concentration while cells are stalled in mitosis has been described previously (43), and are essential for maintaining volume homeostasis. Using quantitative modeling, we demonstrated that exponential growth of cells early in the cell cycle depended on mitochondrial volume/activity in WT cells (Fig. 5C). Our model also predicted linear growth of cells devoid of mitochondrial volume (activity). Therefore, we tested our model in cells lacking mtDNA, and indeed observed that *rho0* cells switched to linear growth, with a constant growth rate across the cell cycle (Fig. 5E, F). Similarly, in Mmb1Δ daughter cell pairs which partitioned the mother cell cytoplasm symmetrically, but mitochondria highly asymmetrically, we observed a marked difference between the growth kinetics of cells that contained little mitochondria and that of the cells that were apportioned more mitochondria (Fig. S5E, F).

Which aspects of mitochondrial function are relevant for the modulation of cell growth kinetics? An intuitive answer would be energy production, since *rho0* cells lack the ability to generate mitochondrial ATP (34). Other studies support this conclusion, since increased mitochondrial mass leads to increased rates of energy-dependent transcription and translation (37, 44). Future studies will also help identify the molecules and machinery that participate in the crosstalk between mitochondrial function and cell growth. While it has been established in several systems that mitochondrial volume and mtDNA scale with cell size (3, 45, 46), here we additionally show that mitochondrial volume is a key determinant of *when* the size control mechanisms are invoked.

In summary, our findings reveal a robust mechanism of mitochondrial homeostasis governed by mitochondrial activity: when cells inherit a lower-than-average mitochondrial volume, they compensate by extending their cell cycle, thereby synthesising additional mitochondria before dividing in accordance with the sizer mechanism. This compensatory strategy preserves mitochondrial homeostasis over multiple generations of dividing cells.

## Materials and Methods

### Growth media and conditions

In this study, the S. pombe cells were grown in yeast extract (YE5) medium with appropriate supplements at a temperature of 30^*°*^C (47).

### Construction of strains

The fission yeast strains used in this study are listed in Supplementary Table S3. Strain VA102 was constructed by crossing PT1650 (h+ cox4GFP:leu1 ade6-M210 ura4-D18; see Table S1) with JCF4627 (hade6-M210 leu1-32 ura4-D18 his3-D1 hht1mRFP-hygMX6). Strain VA125 was constructed by crossing TFSP669 (hura4-D18 tuf1-mRFP::ura4 sdh2-mEGFP::nat) with L975 (h+) and selecting for GFP positive and RFP negative cells. Strain VA127 was constructed by crossing PT1650 with (h+ cox4-GFP:leu1 ade6-M210 ura4-D18) with YSM3340 (h90 ura4+::pact1-mCherry-D4H. Strain VA130 was constructed by crossing JFY3062 (h+ tom20mCherry:NatR) with YSM3811 (hura4+::pact1:sfGFP- 3RitCb:terminatortdh1 ade6-M210 leu+). VA131 was constructed by crossing crossed VA078 (h+ mmb1Δ:Kanr) with VA130 (h- ura4+::pact1:sfGFP-3RitCb:terminatortdh1 Tom20-mCherry:NatR). Strain VA135 was constructed by crossing VA132 (h-ase1::kanMX6 leu1-32 ura4-D18 Tom20- mCherry:NatR) with VA128 (h+ ura4+::pact1:sfGFP- 3RitCb:terminatortdh1 leu-). Strain VA136 was constructed by crossing Dnm1Δ (h+ dnm1::kanr leu1-32ade-(ura+)) with VA130 (h- ura4+::pact1:sfGFP-3RitCb:terminatortdh1 Tom20-mCherry:NatR). Strain VA144 was constructed by crossing FY20823 (leu1 ura4 his7 Δklp5::ura4+ Δklp6::ura4+) with JFY3062 (h+ tom20-mCherry:NatR).

### Preparing the YMM mould

The YMM design used in this study is identical to what was described in Ref. (22). We used an epoxy resin replica mould of the YMM which is a robust and cost-effective alternative to the traditional silicon wafer-based master mould (23). To make the epoxy resin replica mould, first a polydimethylsiloxan (PDMS) template was made from the silicon wafer-based master mould. A thin PDMS layer (10: 1 monomer-to-cross-linker ratio) was cured in a Petri dish. The PDMS template was then plasma-cleaned, placed feature-side up on the cured PDMS layer, and covered with epoxy resin. After being degassed under a vacuum to remove air bubbles, the resin was allowed to fully cure. Finally, the Petri dish was removed using a Dremel tool and the PDMS template was carefully separated from the hardened epoxy. The resulting epoxy mould retained the microchannel pattern and served as a stable and reusable master mould for producing multiple YMM devices.

### Preparing the YMM

A 10:1 PDMS mixture was degassed to remove bubbles that formed due to mixing, poured into the epoxy resin mould, and cured overnight at 65°C; the cured PDMS was peeled off, trimmed, and punched to form inlets/exit holes to allow the insertion of microfluidic tubing. To remove dust particles from the surface of the coverslips, 24×60 mm coverslips (no. 1.5) were soaked overnight in 1% SDS plus 0.1 M NaOH solution, rinsed with MilliQ water, sonicated in isopropanol for a couple of minutes and dried using nitrogen air to prevent liquid stains on the glass surface. Then both coverslips and PDMS devices were plasma-cleaned (feature side up), brought into contact, and placed in a 65°C oven for at least an hour to bond. Leak testing was performed in a custom antileak chamber to ensure a stable, spill-free environment for extended time-lapse imaging.

### Preparing *S. pombe* for imaging in the YMM

To populate the YMM with cells, the device was first wet with 0. 01% Triton and thoroughly rinsed with YE5 medium. A highly concentrated cell suspension (nearly a cell pellet) was then loaded via a short piece of YMM tubing and a syringe fitted with a blunt needle. The cell filled device was centrifuged at 4000-4500 rpm (with rotation by 90°if necessary) until the microchannels were packed (~5 - 10 minutes). Excess cells were removed from the inlet and outlet holes using the short piece of YMM tubing and a syringe fitted with a blunt needle, allowing the outflow to rinse away stray cells without blocking the outlet. A peristaltic pump was then attached to supply fresh YE5 medium and this was input into the YMM using fresh tubes in the inlet and outlet holes. As cells grew in lines within the YMM, any cells that were pushed out of the growth channels were washed away via the feeding channel.

### Live-cell imaging using the YMM

To image *S. pombe* cells in the YMM, they were first acclimated by running the setup overnight on a Nikon confocal microscope (A1 or AX- R2) using the built-in perfect focus system (PFS) feature to maintain focus. Faster flow rates (≥ 2 ml/h) helped align cells with the original cell (mother) trapped at the closed end of the channel while the progeny (daughters) escaped as the mother and daughters divided. Proper connections between the peristaltic pump and YMM tubing (via a needle and male luer) prevented leaks. Sufficient medium was provided to ensure continuous flow and cell viability. For strains lacking RitC:GFP, to delineate cell boundaries (PHP14 and VA144, see Supplementary Table S3), YE5 media supplemented with 2*µ*g/mL calcofluor white stain (Sigma-Aldrich cat. no.:18909) was continuously flowed into the YMM. The peristaltic tubing was replaced after each experiment and the flow rates of the pump recalibrated as needed. Confocal microscopy (Nikon A1 or AX-R2, 60× water-immersion objective with 1.20 NA) was used to acquire z-stacks (1 *µ*m intervals) and time-lapse images (5–15 min intervals over 30–60 h) in transmitted light, 488 and 561 nm channels, captured both mitochondria (Tom20:mCherry) and cell boundaries (RitC:GFP).

### Image and data analysis

We developed an automated image analysis framework for segmenting and tracking *S. pombe* growing in the YMM using Fiji/ImageJ (48), MATLAB (Mathworks Corp.), and Python. First, the fluorescent images of the YMM were cropped into individual channels and each channel was oriented with the closed end, containing the mother cell, at the bottom. Images with the closed end at the top were rotated 180° to maintain consistency. The images with fluorescent cell membranes were denoised with Nikon’s AI-driven algorithm to remove Poisson shot noise and enhance the signal-to-noise ratio. The denoised images were then segmented using Cellpose, which is a deep learning tool trained on various cell morphologies (49). Cells were segmented using either ‘cyto2’, ‘cyto3’, or a custom Cellpose model. In instances where the default cyto models produced segmentation errors on Calcofluor-stained images, a custom model was trained on the data to ensure segmentation accuracy. During segmentation, the diameter parameter was set to ‘None’, allowing the algorithm to automatically estimate the optimal cell size based on image characteristics. After segmentation, the ‘regionprops’ function in MATLAB extracted quantitative cell features (e.g. centroid, bounding box, length, width, and area).

We define cell birth as the moment the membrane (RitC:sfGFP) signal first appears at the medial plane of the cell, marking the onset of septum formation. This point also coincides with cell segmentation by Cellpose. In *S. pombe*, the septum, composed of specific polysaccharides, acts as a new cell wall that divides the two daughter cells (48, 50, 51). During cytokinesis, septum formation physically separates the cytoplasm, enabling each daughter cell to grow independently (52, 53). Following septation, each daughter cell briefly passes through the G1 phase before entering the S phase to initiate its own DNA replication cycle (54).

The cells were arranged by the y-coordinate of their centroids to identify and track the bottom-positioned mother cells. In the tracking algorithm, a decrease exceeding 30% in the area of a mother cell between consecutive frames was interpreted as a division event, which led the algorithm to visually mark the mother and daughter cells with bounding boxes. Each cell received a unique identifier (ID) recording its generation and lineage (e.g. mother cell ‘001’ producing daughters ‘011’ and ‘012’, see Supplementary Video S1). Mother cells were tracked continuously throughout the movie, while daughter cells were tracked until they underwent division. Finally, the areas/lengths of the mother and daughter cells were plotted, along with their mitochondrial intensities or volumes over time.

The mitochondrial channel was binarized using 3D Trainable Weka Segmentation in a two-class system, with mitochondria defined as one class and the background as the other (55). For each field of view (FOV), a single representative 3D image was manually classified to construct a classifier model, which was then applied to all images in that time-lapse. Owing to differences in mean intensities between FOVs, a distinct classifier model was developed for each field. The mitochondrial area for each z-slice was determined using MATLAB’s built-in function ‘bwarea’, which estimates the area of the objects in the binary image and then scaled by the corresponding pixel value. The mitochondrial volume was calculated by summing these area values across all z-slices and multiplying the sum by the inter-slice distance. For quantifying mitochondrial area and intensity, sum intensity projection images were used. The mitochondrial area was again extracted using ‘bwarea’ and scaled by the pixel value, while integrated density (which is the sum of all pixel intensities within the region), was measured to determine mitochondrial intensity. To further ensure uniformity, all mitochondrial volume and intensity values were normalised to the mean mitochondrial volume/intensity for each FOV.

### Measurement of surface area and volume of *S. pombe* cells

The surface area of the cell was calculated as follows:

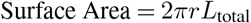

where *r* is the radius of the cell, calculated as half of the width of the cell, *L*_total_ is the total measured cell length, including its hemispherical ends and *π* is the mathematical constant.

The volume of the cell was calculated as follows:

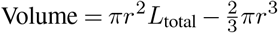

### Plotting and statistics

All plots were generated in MATLAB. For population-wide patterns (Fig.1C and D, Fig.2B and C, Fig.3A, B, C, E and F, Fig.S2 C, D and E, Fig. S3, S4A), raw values were *rescaled* by normalising to their respective means (e.g., in Fig.2C, the birth lengths were divided by the mean birth length, and the mitochondrial volumes at birth were divided by the mean mitochondrial volume at birth). In Fig.3D, the normalised mitochondrial volume refers to mitochondrial volume normalised by FOV. In Fig. 4C, the cell length of each daughter was normalised to the sum of the daughter lengths. In Fig. 5, the cell area data was normalised to the mean birth areas and the mitochondrial volumes to the mean volume per FOV and the cells included in the plots have cell cycle durations within 1 SD of the mean duration of the WT cell cycle. In Fig. S5, the mitochondrial volume was normalised to the mean mitochondrial volume per FOV.

Heat scatter plots were created using a custom MATLAB function (56) to depict local data density with a colour map (warmer colours indicate higher density). Data were grouped into bins, and outliers were removed using MATLAB’s ‘rmoutlier’ function. Trend lines were fit using MATLAB’s ‘cftool’ with a Linear Model (‘Poly1’) and each point weighted by the inverse of its SEM. Shaded error plots, showing the mean with SEM or SD shading, were generated using the custom ‘shadedErrorBar’ function (57), with similar fitting procedures. For all the shaded error bar plots involving the mutants, 5 bins were used. Data were checked for normality using the ‘chi2gof’ function in Matlab. Then, to test the statistical significance of the difference between distributions we used ordinary one-way ANOVA or Student’s T-test for parametric data and Kruskal-Wallis test or Mann-Whitney test for non-parametric data. The figures were organised and prepared in Illustrator.

## Acknowledgements

We thank the Katharina Gaus Light Microscopy Facility, UNSW, Australia for the use of the Nikon A1, AX-R2 confocal microscopes and data analysis computer; P. Delivani (Max Planck Institute of Molecular Cell Biology and Genetics, Dresden, Germany), S. Martin (University of Geneva, Switzerland), Tomoyuki Fukuda (Niigata University Graduate School of Medical and Dental Sciences: Niigata, Japan), J. Friedman (UT Southwestern Medical Center, USA), M. Takaine (Gunma University, Gunma, Japan), I. Tolić (Rudēr Bošković Institute, Zagreb, Croatia), P. Tran (University of Pennsylvania, Philadelphia, PA, USA), T.D. Fox (Cornell University, Ithaca, NY, USA), and National BioResource Project Japan for yeast strains and constructs, EMBL Australia for funding to VA and RM, Australian Research Council Centre of Excellence for Mathematical Analysis of Cellular Systems (CE230100001) for funding to RM, and NIH grant R35 GM130327 for funding to WM.

## Notes

### Competing Interest Statement

The authors have declared no competing interest.

