## Supplementary Information for "Mitochondrial function regulates cell growth kinetics to actively maintain mitochondrial homeostasis"

This document contains 5 Supplementary Figures, 3 Supplementary Tables, and 3 Supplementary Video Captions.

### Supplementary Figures

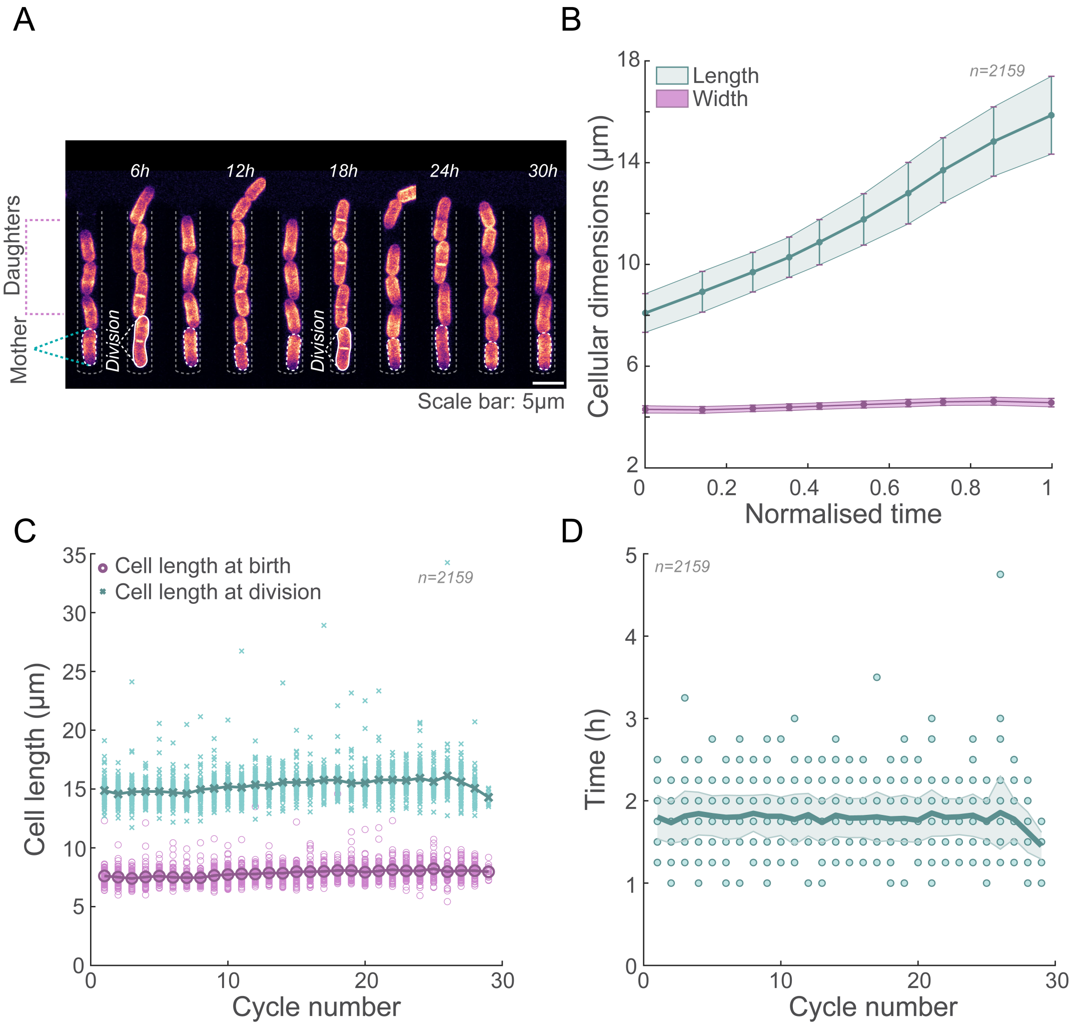


**Figure S1. Cell growth is optimal in the YMM**. **(A)** A representative montage of cells expressing a fluorescent membrane marker (RitC:GFP) growing in a single channel of the YMM, showing representative mother and daughter cells, as well as division events. **(B)** Plot depicting changes in cell length and width within the YMM during the cell cycle, **(C)** Plot of the raw (scatter) and mean (solid line) cell lengths of WT cells growing in the YMM at birth (magenta) and at division (teal), **(D)** Plot of the individual cell cycle durations (circles) and mean ± SD (solid line and shaded region respectively) of WT cells growing in the YMM. Strain VA130 was used in this figure, see Supplementary Table S3.

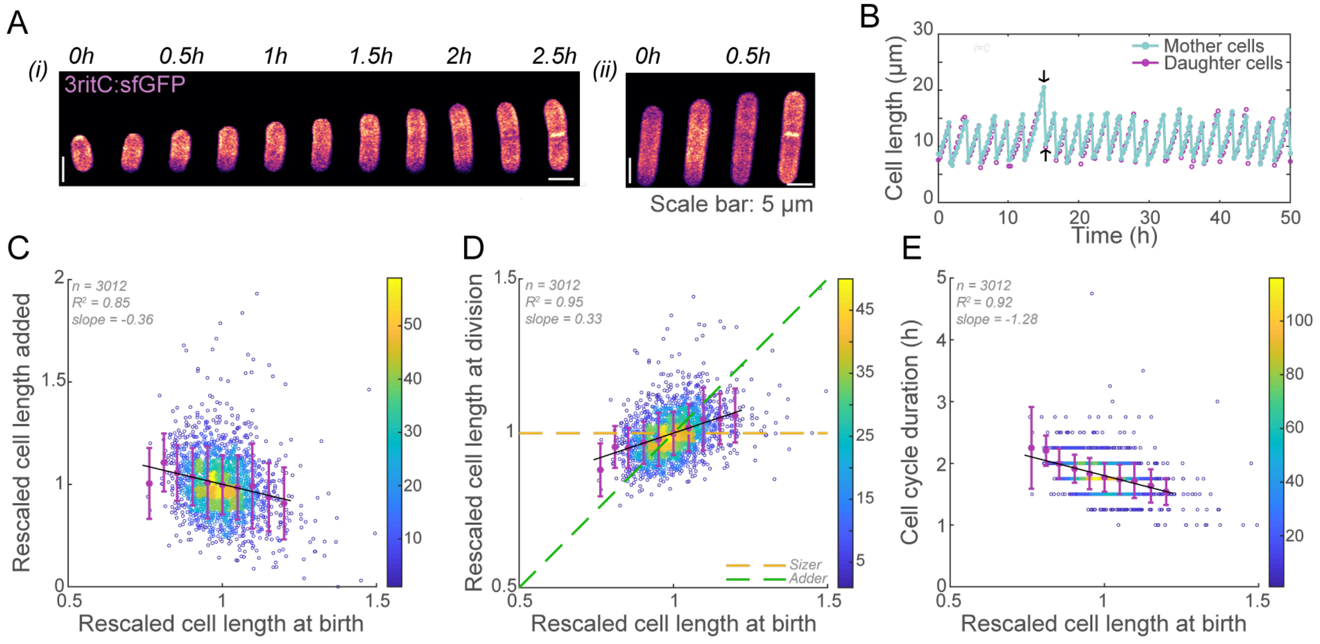


**Figure S2. *Schizosaccharomyces pombe* cells show "sizer-like" behaviour**. **(A)** Example montages of an entire cycle of cells expressing a fluorescent membrane marker with different sizes at birth (i - small cell and ii - large cell). **(B)** Example plot of cell length over time in WT cells growing in the YMM showing cell length fluctuations. The arrow heads point to a cell with increased length at division (top arrowhead) and birth (bottom arrowhead), **(C)** Cell length added during the cycle plotted against the cell length at birth shows an inverse relationship, **(D)** Cell length at division plotted against the cell length at birth shows a sizer-like behaviour. The yellow dashed line represents the expected data for a perfect sizer and the green dashed that for an adder, **(E)** Cell length at birth plotted against the cell cycle duration shows that cells that are smaller at birth spend more time in the cell cycle. In **C-E**, error bars represent the SD and the heat map represents the density of the data. Strain VA130 was used for all the figures in this panel (see Supplementary Table S3).

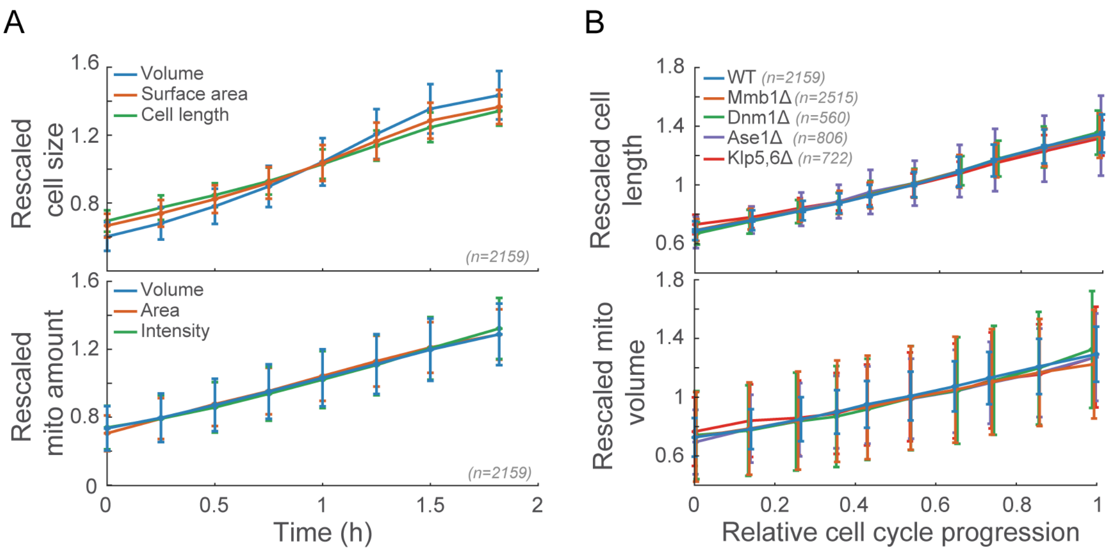


**Figure S3. The mitochondrial amount grows with cell size**. **(A)** Plots of cell size (cell volume, surface area and length) over time (top) and the mitochondrial amount (mitochondrial volume, area and intensity) over time (bottom). The error bars represent the mean $\pm$ SD. The volume and surface area of the cell were calculated as detailed in the Methods section. Strain VA130 was used for this panel (see Table S1). **(B)** Plots of cell length over time (top) and the mitochondrial volume over time (bottom). The solid lines represent the mean and the error bars SD. The data in this figure were rescaled as detailed in the Methods section. Strains VA130, VA131, VA136, VA135, and VA144 were used for for this panel (see Supplementary Table S3).

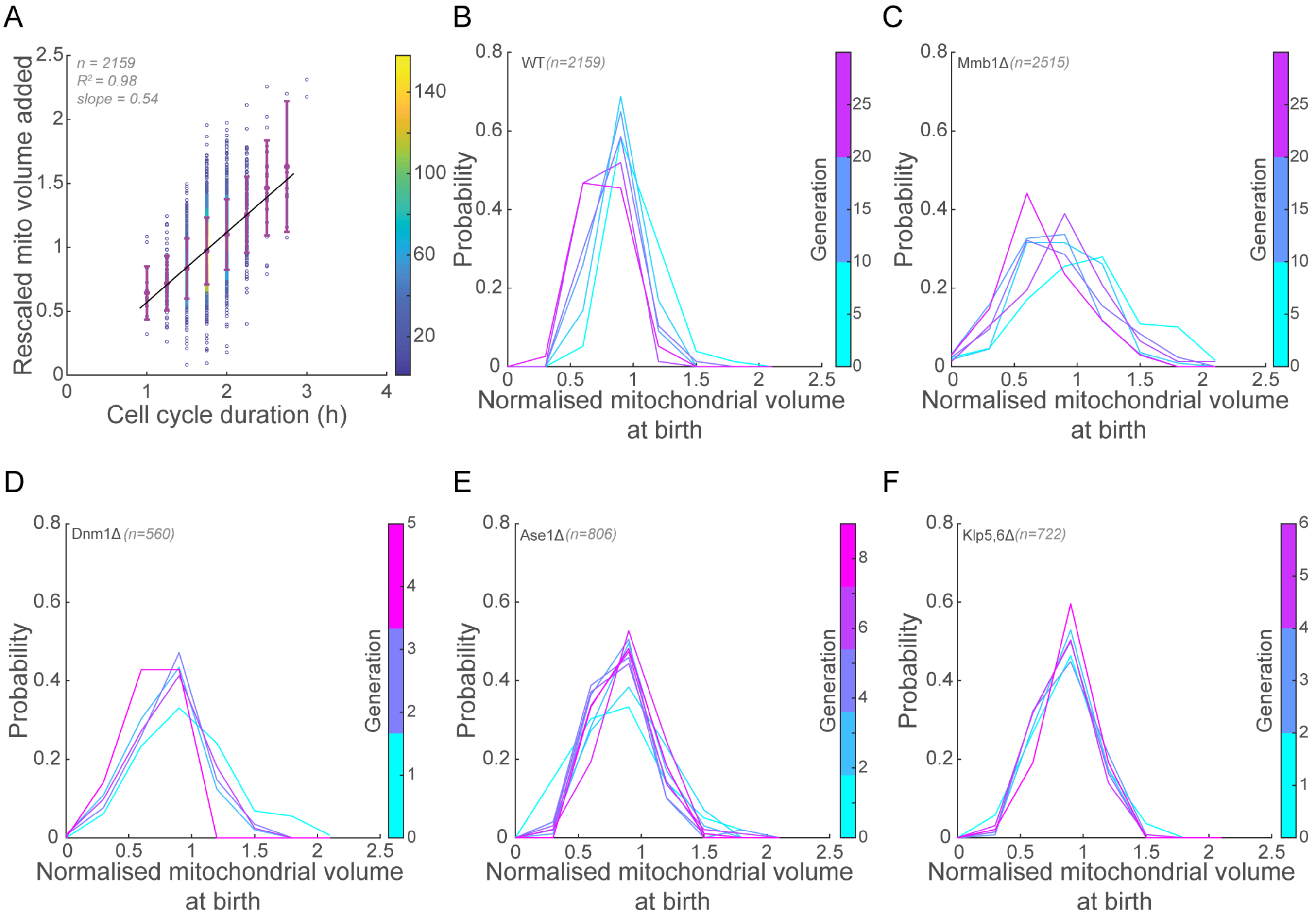


**Figure S4. WT and mutant cells correct for errors in mitochondrial partitioning within a single generation A** The mitochondrial volume added per cycle and the duration of cell cycle in WT cells show a linear relationship, i.e., longer the cell cycle, more the mitochondrial volume added before division. Error bars represent SD. Black solid lines represent linear fits to the data; the heat map represents the density of the data **(B)** Histogram of mitochondrial volume at birth for multiple generations of WT, **(C)** Mmb1△, **(D)** Dnm1△, **(E)** Ase1△, and **(F)** Klp5,6△ cells tracked in the YMM. The data in **B** and **C** were plotted for every fifth generation. Strains VA130, VA131, VA136, VA135, and VA144 were used for for this panel (see Supplementary Table S3)

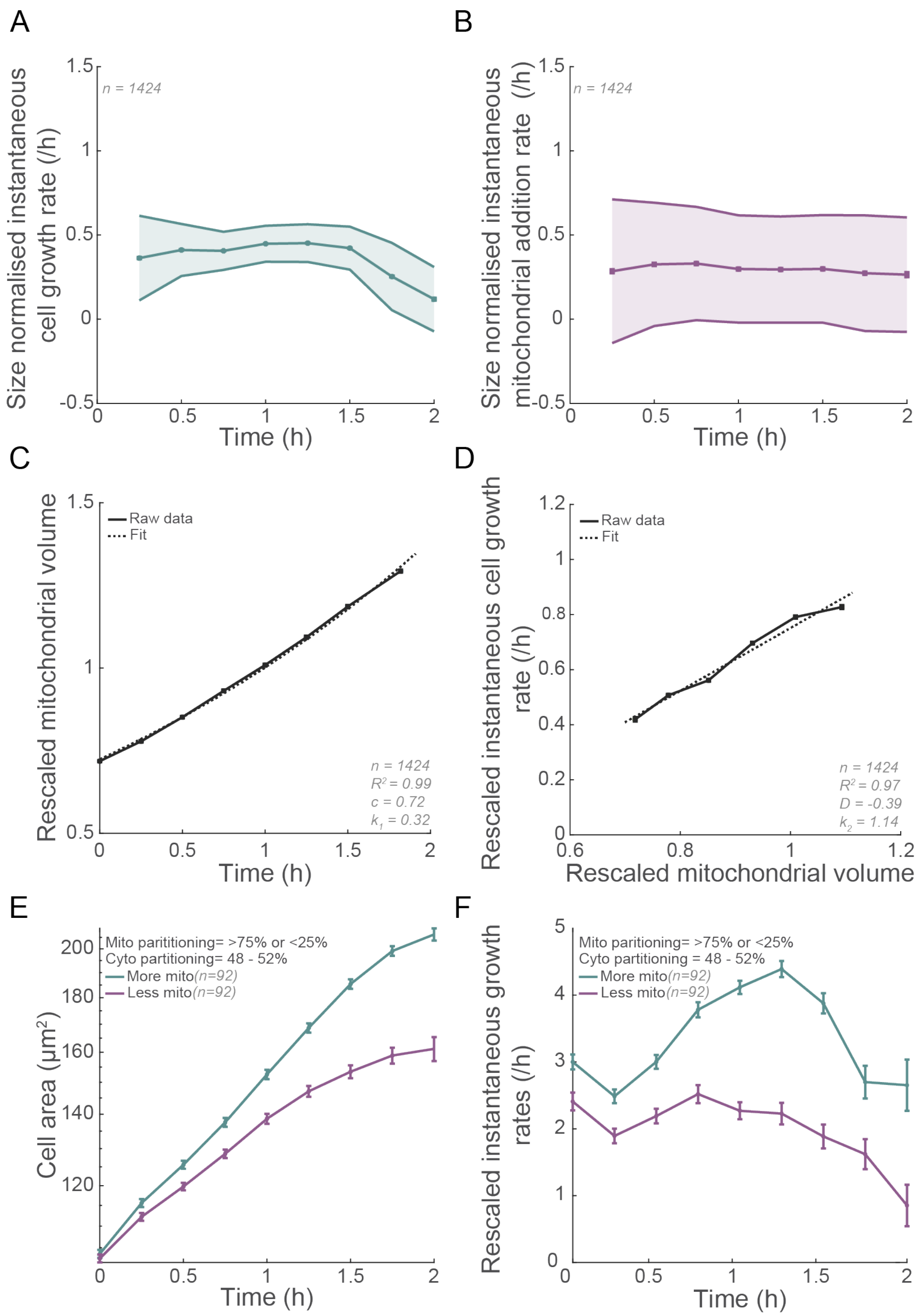


**Figure S5. Mitochondrial activity leads to exponential growth of cells (A)** Plot of size normalised instantaneous growth rate of WT cells ($\frac{1}{A}\frac{dA}{dT}$) over time, **(B)** Plot of volume normalised instantaneous mitochondrial addition rate ($\frac{1}{M}\frac{dM}{dT}$) over time in WT cells. In A and B, the solid line represents the mean and the shaded region represents the SD, **(C)** Plot of mitochondrial volume over time fit to an exponential curve, **(D)** Plot of the instantaneous growth rates of the cell against the mitochondrial volume. The solid line represents the raw data and the dashed line represents the fit to a polynomial of degree 1. **(E)** Plot of cell area over time for Mmb1$\Delta$ cells that divided symmetrically but partitioned mitochondria asymmetrically, showing a marked switch of cells that inherited <25% of the mother mitochondria to linear growth, **(F)** Plot of instantaneous growth rate of cells over time for Mmb1$\Delta$ cells that divided symmetrically but partitioned mitochondria asymmetrically, demonstrating nearly constant growth rates in cells that inherited <25% of the mother mitochondria. The data in this figure were rescaled as detailed in the Methods section. Strains VA130 and VA131 were used for for this panel (see Supplementary Table S3)

#### Supplementary Table S1: Measured parameters for WT and mutant cells

| **Strain** | **Cell length at birth** (mean ± SD) $\mu$m | **Cell length at division** (mean ± SD) $\mu$m | **Cell cycle duration** (mean ± SD) h |
|---|---|---|---|
| WT | 8.1 ± 0.8 | 15.8 ± 1.4 | 1.8 ± 0.3 |
| Mmb1 | 7.9 ± 0.8 | 15.3 ± 1.4 | 2.0 ± 1.9 |
| Dnm1 | 7.4 ± 0.9 | 14.9 ± 1.6 | 2.0 ± 0.4 |
| Ase1 | 7.6 ± 1.1 | 14.8 ± 2.4 | 1.9 ± 0.4 |
| Klp56 | 7.8 ± 0.7 | 14.1 ± 1.2 | 1.8 ± 0.3 |
| *rho0* | 9.8 ± 1.4 | 18.5 ± 2.9 | 9.3 ± 4.9 |

#### Supplementary Table S2: Goodness of fits ($R^2$) and slopes for mutant data

| **Strain** | $R^2$ | **Slope** |
|---|---|---|
| **Fig. 1C** | | |
| WT | 0.98 | -1.47 |
| Mmb1Δ | 0.99 | -1.87 |
| Dnm1Δ | 0.98 | -1.83 |
| Ase1Δ | 0.93 | -2.77 |
| Klp5,6Δ | 0.99 | -1.39 |
| **Fig. 1D** | | |
| WT | 0.99 | 0.31 |
| Mmb1Δ | $6.4 \times 10^{-5}$ | 0 |
| Dnm1Δ | 0.75 | 0.07 |
| Ase1Δ | 0.14 | -0.05 |
| Klp5,6Δ | 0.98 | 0.31 |
| **Fig. 2C** | | |
| WT | 0.99 | 0.68 |
| Mmb1Δ | 0.99 | 0.93 |
| Dnm1Δ | 0.98 | 0.41 |
| Ase1Δ | 0.98 | 1.08 |
| Klp5,6Δ | 0.92 | 0.74 |
| **Fig. 3E** | | |
| WT | 0.99 | -0.49 |
| Mmb1Δ | 0.99 | -0.84 |
| Dnm1Δ | 0.69 | -0.30 |
| Ase1Δ | 0.93 | -0.50 |
| Klp5,6Δ | 0.98 | -0.97 |
| **Fig. 3F** | | |
| WT | 0.91 | 0.16 |
| Mmb1Δ | 0.98 | 0.19 |
| Dnm1Δ | 0.89 | 0.05 |
| Ase1Δ | 0.92 | 0.32 |
| Klp5,6Δ | 0.03 | 0.03 |

#### Supplementary Table S3: List of strains used in this study

| Name | Genotype | Source |
|---|---|---|
| Dnm1Δ | h- dnm1::kanr leu1-32ade- | Yannick Gachet, Toulouse |
| FY20823 | h- leu1 ura4 his7 Δklp5::ura4+ Δklp6::ura4+ | YGRC, Japan |
| G3B | h- Δ klp5- Δ Δklp6 – nmt1-GFP-atb2 leu ade | Rafael Carazo Salas, UK |
| HN1007 | h- SPBC1348.11«HphMX6-Padh1-mCherry | Hidenori Nakaoka, The University of Tokushima, Japan |
| JCF4627 | h- ade6-M210 leu1-32 Hht1-mRFP:kanMX6::leu1 | Julia Promisel Cooper, University of Colorado, USA |
| JFY3062 | h- tom20-mCherry:NatR | Jonathan Friedman, UT Southwestern, Dallas, Texas, USA |
| L975 | h+ WT | Iva Tolic' |
| MMY3246 | h? leu1-32 ura4-D18 aco1:GFP:ura4+ | Fuyuki Ishikawa, Koyoto University, Japan |
| PHP14 | h- rho0 ade6M-216 leu1-32 ptp1-1 | Thomas Fox, Cornell University, USA |
| PT1650 | h+ cox4-GFP:leu1 ade6-M210 ura4-D18 | Phong Tran, USA |
| PT2244 | h+ mmb1Δ:Kanr cox4-GFP:leu2 mCherry- atb2:Hygr ade6-m210 leu1-32 ura4-d18 | Phong Tran, USA |
| PT592 | h- ase1::kanMX6 leu1-32 ura4-D18 | Phong Tran, USA |
| TFSP669 | h- ura4-D18 tuf1-mRFP::ura4 sdh2-mEGFP::nat | Tomoyuki Fukuda, Niigata University, Japan |
| VA078 | h+ mmb1Δ:Kanr | This study |
| VA102 | h- hht1-mRFP-hygMX6 cox4-GFP:leu1 ade6-M210 leu1-32 ura4-D18 | This study |
| VA125 | h90 sdh2-mEGFP::nat | This study |
| VA127 | h? ura4+::pact1-mCherry-D4H cox4-GFP:leu1 | This study |
| VA128 | h+ ura4+::pact1:sfGFP-3RitCb:terminatortdh1 leu- | This study |
| VA130 | h- ura4+::pact1:sfGFP-3RitCb:terminatortdh1 Tom20-mCherry:NatR | This study |
| VA131 | h+ mmb1Δ:Kanr ura4+::pact1:sfGFP-3RitCb:terminatortdh1 Tom20-mCherry:NatR | This study |
| VA132 | h- ase1::kanMX6 leu1-32 ura4-D18 Tom20-mCherry:NatR | This study |
| VA135 | h- ase1::kanMX6 Tom20-mCherry:NatR ura4+::pact1:sfGFP-3RitCb:terminatortdh1 | This study |
| VA136 | h+ dnm1::kanr ura4+::pact1:sfGFP-3RitCb:terminatortdh1 Tom20-mCherry:NatR | This study |
| VA144 | h- Δklp5::ura4+ Δklp6::ura4+ Tom20-mCherry:NatR | This study |
| YSM3440 | h90 ura4+::pact1-mCherry-D4H | Sophie Martin, University of Geneva, Switzerland |
| YSM3811 | h- ura4+::pact1:sfGFP-3RitCb:terminatortdh1 ade6-M210 leu+ | Sophie Martin, University of Geneva, Switzerland |

#### Supplementary Video Captions

**Video S1. *S. pombe* cells growing in the YMM for 60 h**
Video depicting a single channel of the YMM populated with *S. pombe* cells whose cell boundaries are tagged with D4H:mCherry (grey, strain YSM3440, see Supplementary Table S3). The mother and daughter cells are represented by the teal and magenta ROIs. The cell IDs for the mother and daughter cells are depicted beside their ROIs. The time interval between frames is 15 min, z-slices are 1 $\mu m$ apart for a total of 6 $\mu m$. The scale bar represents 10 $\mu$m, time is indicated in hh:mm.

**Video S2. Ase1$\Delta$ cells growing in the YMM for 10 h**
Video depicting a single channel of the YMM populated with Ase1$\Delta$ cells whose cell boundaries are tagged with RitC:GFP (grey, strain VA135, see Supplementary Table S3). The mother and daughter cells are represented by the teal and magenta ROIs. The cell IDs for the mother and daughter cells are depicted beside their ROIs. The time interval between frames is 15 min, z-slices are 1 $\mu m$ apart for a total of 6 $\mu m$. The scale bar represents 10 $\mu$m, time is indicated in hh:mm.

**Video S3. *S. pombe* cells tagged with Tom20:mCherry growing in the YMM**
Video depicting five growth channels of the YMM populated with *S. pombe* cells whose mitochondria are tagged with Tom20:mCherry (green) and cell boundaries are tagged with RitC:GFP (magenta, strain VA130, see Supplementary Table S1). The time interval between frames is 15 min, z-slices are 1 $\mu$m apart for a total of 6 $\mu$m. The scale bar represents 10 $\mu$m, , time is indicated in hh:mm.
